# INDIVIDUAL LIGHT HISTORY MATTERS TO COPE WITH THE ANTARCTIC SUMMER

**DOI:** 10.1101/2022.12.29.522237

**Authors:** Julieta Castillo, André Comiran Tonon, María Paz Loayza Hidalgo, Ana Silva, Bettina Tassino

## Abstract

The effect of light, main zeitgeber of the circadian system, depends on the time of day it is received. A brief trip to the Antarctic summer (ANT) allowed us to explore the impact of a sudden and synchronized increase in light exposure on activity-rest rhythms and sleep patterns of 11 Uruguayan university students, and to assess the significance of light history in determining individual circadian phase shift. Measurements collected in the peri-equinox in Montevideo, Uruguay (baseline situation, MVD) and in ANT, included sleep logs, actigraphy, and salivary melatonin to determine dim-light melatonin onset (DLMO), the most reliable marker of circadian phase. The increase in light exposure in ANT with respect to MVD (affecting both light-sensitive windows with opposite effects on the circadian phase) resulted in no net change in DLMO among participants as some participants advanced their DLMO and some others delayed it. The ultimate cause of each participant’s distinctive circadian phase shift relied on the unique change in light exposure each individual was subjected to between their MVD and ANT. This is the first study to show a clear physiological effect of light either advancing or delaying the circadian phase dependent on individual light history in an ecological study.

## INTRODUCTION

The exposure to the light–dark cycle is the most powerful environmental synchronizing signal for the circadian system across evolution. Light impacts on specific retinal ganglion cells, whose axons reach the mammalian circadian pacemaker, the suprachiasmatic nuclei, located in the hypothalamus^1,2^. The effect of light on the circadian phase depends on the moment of the day in which light exposure occurs, following a well-documented phase response curve first described in rodents^3^, but also confirmed in humans^4–7^. Light exposure occurring in the evening (early biological night) induces a phase-delay shift of the circadian pacemaker, whereas light exposure occurring early in the morning (late biological night) induces a phase-advance shift. Modern urban life has disrupted the light patterns expected by the biological clock, leading to changes that are difficult to predict^8–10^ and that are highly variable among individuals^11^. However, it seems clear that all the light and darkness to which humans are exposed throughout the day plays a role in entrainment^12^. As melatonin is inhibited by light, the detection of the onset of the nocturnal increase of melatonin (i.e., the dim light melatonin onset, DLMO) has emerged as the best predictor of the beginning of the endogenous dark phase^13,14^, which also accounts for the reported light-dependent changes and individual variability^11^.

Although distorted by electric light and variable across latitudes and seasons, light exposure still impacts on the human circadian system in normal life, leading to light-dependent phase-shifts and changes in sleep patterns. For example, around the equator, rubber tappers exposed to electric light at home, and therefore more exposed to evening light, showed later DLMO and sleep than workers without electric light at home^15^. On the other hand, a seasonal experiment carried out in natural camping conditions in Colorado, USA (∼40°N, with a seasonal light-dark change of around 10/14 to 14/10) reported the expected longer nocturnal melatonin pulse in winter than in summer, but no significant changes in DLMO across seasons^16^, maybe because the longer sunlight of summertime impacts in both the phase-advance and phase-delay windows. When seasonal changes in photoperiod are more extreme, for example in Antarctic scientific bases (∼75°S, with a seasonal light-dark change of around 0/24 to 24/0), circadian rhythms during winter, including melatonin, are usually delayed or even completely desynchronized, adopting free-running patterns^17,18^. Moreover, in Antarctic^17^ as well as in Arctic populations^19^, sleep is also delayed and more disturbed in winter, reinforcing the importance of the synchronizing power of morning summer light. These ecological human chronobiological experiments evaluating the effects of light in normal life show the complexity of the light entrainment of the circadian system and make it difficult to extract general rules of its impact on circadian rhythms and sleep patterns. Abrupt changes in lighting conditions offer a different angle of analysis that reveals the plasticity of the circadian system to cope with short-term changes. The iconic camp experiment^20^ that evaluated the effect of a sudden change in light exposure (electric light vs natural light) between two successive weeks showed a re-set of the circadian phase in natural light conditions; i.e., after only seven days of natural lighting, DLMO synchronizes with sunset and among individuals. In addition to this remarkable plasticity of the circadian system to adjust to short-term changes, recent studies have focused on the differences in individual responses to similar light exposure profiles^21–23^. Overall, the effects of light on sleep and circadian phase depend on a wide range of individual traits such as age^24–26^, race/ethnicity^27–29^, sex^30,31^, chronotype^20,32,33^, photosensitivity^11,26^, and photic history^34–36^. Only few studies address the effect of light history on circadian system and sleep in real-world settings, and two of them have taken advantage of Antarctica as a natural laboratory^37,38^. Although these and other investigations confirm that individual photic history (over days and weeks) affects the phase shift in response to light^34,39^, several aspects of this relationship remain unclear, especially in ecological conditions^23^.

Seasonal changes imply gradual variations in photoperiod that involve more light exposure on both the phase-advance and the phase-delay windows. This means an ambivalent signal for the circadian system that forces a trade-off between advancing or delaying the circadian phase with no predictable net effect. Travelling along the same meridian emerges as an interesting ecological model to mimic acute seasonal changes, when occurring in proper times of the year (from winter to summer or vice versa). Theoretical estimations indicate that these trans-latitudinal trips can still have an impact on the circadian system^40^, whose effects on the circadian phase have not been addressed so far. We studied a population of Uruguayan University students, whose average circadian phase did not change in the Antarctic trip (ANT) when compared to their normal life in the fall equinox of Montevideo, Uruguay (MVD), despite the huge increase in light exposure the Antarctic summer implied^33^. This is an exceptional ecological experiment of a trans-latitudinal trip (within the same time zone) that allowed us to explore individual strategies to cope with light increases impacting at the same time both circadian phase-advance and phase-delay windows. To do this, we first provide a comprehensive view of the effects of the ANT on light exposure, activity-rest rhythms, and sleep patterns using objective measurements and self-reports. We then evaluated the importance of light history in shaping the valence (advance or delay) and the magnitude of the individual circadian phase shift.

## RESULTS

In this study we present a longitudinal comparative analysis of the activity-rest and light exposure patterns of 11 healthy University students (8 females, 3 males, 22.45 ± 1.13 years old from Universidad de la República, Uruguay) between the 2016 fall peri-equinox in Montevideo (baseline situation, MVD) and the 2016 summer in Antarctica (Antarctic trip, ANT, Fig. 1). The assessment of chronotype was performed in MVD with data obtained from two largely validated questionnaires: Munich Chronotype Questionnaire (MCTQ)^41,42^ and the Morningness - Eveningness Questionnaire (MEQ)^43,44^. The mid-sleep point on free days corrected for sleep debt on workdays (MSFsc) was, on average, 06:02 ± 0:50 and ranged from 04:40 to 07:35 and the average value of the MEQ scores was 46.45 ± 8.17 and ranged from 32 to 55. The outputs of these questionnaires were significantly correlated (R = -0.69, *p* = 0.018).

**Figure 1.**
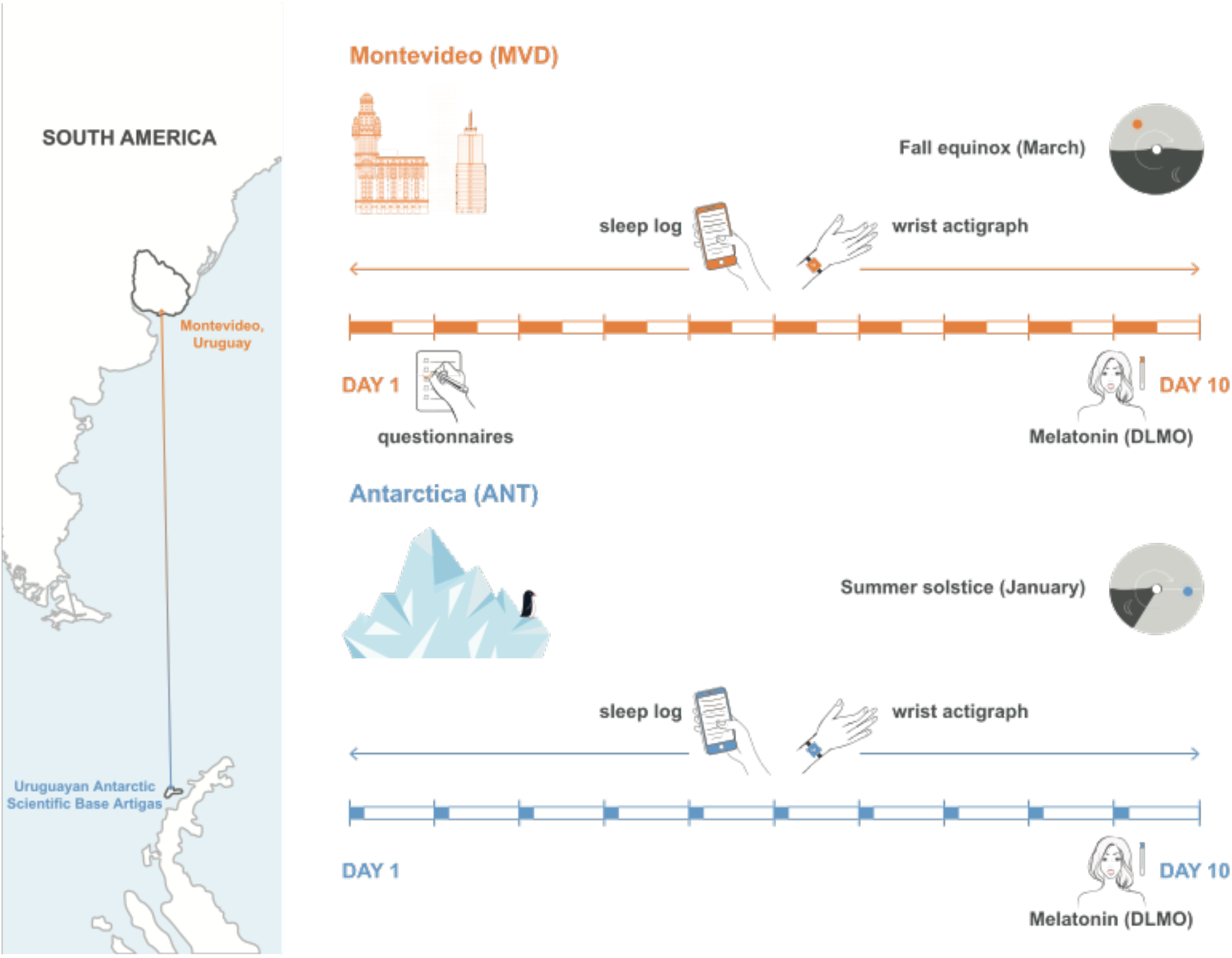
Experimental design and layout. Participants were studied in two conditions: Montevideo (MVD, baseline situation) and during Antarctica trip (ANT), where they filled sleep logs and wore wrist actigraphs continuously for 10 days. On the last day of both locations hourly samples of saliva were collected to determine the dim light melatonin onset (DLMO). In MVD participants also filled questionnaires to assessed chronotypes.

### The impact of the Antarctic summer on the rhythms of activity, light exposure, and sleep

Both conditions, MVD and ANT, represented a similar academic challenge for the participants as MVD corresponded to the start of the university semester in the 2016 austral fall and ANT corresponded to the II Uruguayan Summer Antarctic School held during the 2016 austral summer in the Uruguayan Antarctic Scientific Base Artigas^33,45^. In contrast, light exposure was longer (more minutes exposed to > 1000 lux, Table 1) and more intense (average light exposure of the brightest 10h period, MB10,Table 1) in ANT than in MVD in both the morning and the evening (Fig. 2a). In addition, all participants showed circadian rhythms of light exposure (i.e., with significant fit to a cosine function over 24 h period) in both MVD and ANT. The acrophase of this light circadian rhythm was earlier in ANT than in MVD, while MB10c (midpoint of the brightest 10h period) and LB5c (midpoint of the least bright 5h period) were not affected (Table 1).

**Table 1:** Actigraphy and sleep log data taken on workdays for 8 days in Montevideo and 10 days in Antarctica for all participants (n = 11). Light variables obtained from actigraphy: moment of the day in which the cosinor function reaches its maximum light exposure (light acrophase), midpoint of the brightest 10 h period (MB10c), midpoint of the least bright 5 h period (LB5c), average light exposure of the brightest 10 h period (MB10), average light exposure of the least bright 5 h period (LB5) and the daily average exposure to light intensity above 1000 lux. Activity variables obtained from actigraphy: moment of the day in which the cosinor function reaches its maximum physical activity (acrophase), midpoint of the most active 10 h period (M10c), midpoint of the least active 5 h period (L5c), average activity of the most active 10 h period (M10), average activity of the least active 5 h period (L5), relative amplitude (RA) and the daily average of time participants spent in moderate-vigorous physical activity (MVPA).The subjective sleep quality was self-reported in a sleep-log by a visual analogue scale (VAS). Statistically significant tests (p < 0.05) are shown in bold.

|  | Montevideo | Antarctica | p-value |
| --- | --- | --- | --- |
| **Light variables (Actigraphy)** | Median ± MAD | | Wilcoxon signed-rank test |
| Light acrophase (hh:mm) | 13:33 ± 00:29 | 12:59 ± 00:06 | **0.001** |
| MB10c (hh:mm) | 14:01 ± 00:36 | 13:54 ± 00:05 | 0.916 |
| LB5c (hh:mm) | 04:33 ± 00:46 | 04:10 ± 00:19 | 0.354 |
| MB10 (lux) | 1370.40 ± 215.77 | 4342.63 ± 342.33 | **0.001** |
| LB5 (lux) | 0.042 ± 0.04 | 0.13 ± 0.13 | 0.447 |
| Light exposure >1000 lux (min) | 150.32 ± 44.63 | 223.55 ± 9.44 | **0.003** |
| **Activity variables (Actigraphy)** | Median ± MAD | | Wilcoxon signed-rank test |
| Acrophase (hh:mm) | 16:47 ± 00:22 | 14:26 ± 00:11 | **0.001** |
| M10c (hh:mm) | 16:47 ± 00:34 | 12:50 ± 00:09 | **0.001** |
| L5c (hh:mm) | 05:24 ± 00:32 | 04:22 ± 00:17 | 0.127 |
| M10 (g*min) | 40.72 ± 6.55 | 54.02 ± 5.57 | **0.002** |
| L5 (g*min) | 8.08 ± 1.23 | 7.46 ± 1.30 | **0.005** |
| RA | 0.67 ± 0.08 | 0.75 ± 0.015 | **0.001** |
| MVPA >100 mg (min) | 168.21 ± 32.21 | 283.22 ± 34.31 | **0.001** |
| **Sleep variables (Sleep Logs)** | Mean ± SD | | t test |
| Midsleep point (hh:mm) | 05:08 ± 00:55 | 04:21 ± 00:12 | **0.033** |
| Sleep duration (h) | 7.45 ± 0.85 | 6.08 ± 0.39 | **0.001** |
| Subjective sleep quality (VAS) | 6.76 ± 1.47 | 6.83 ± 1.03 | 0.882 |

**Figure 2.**
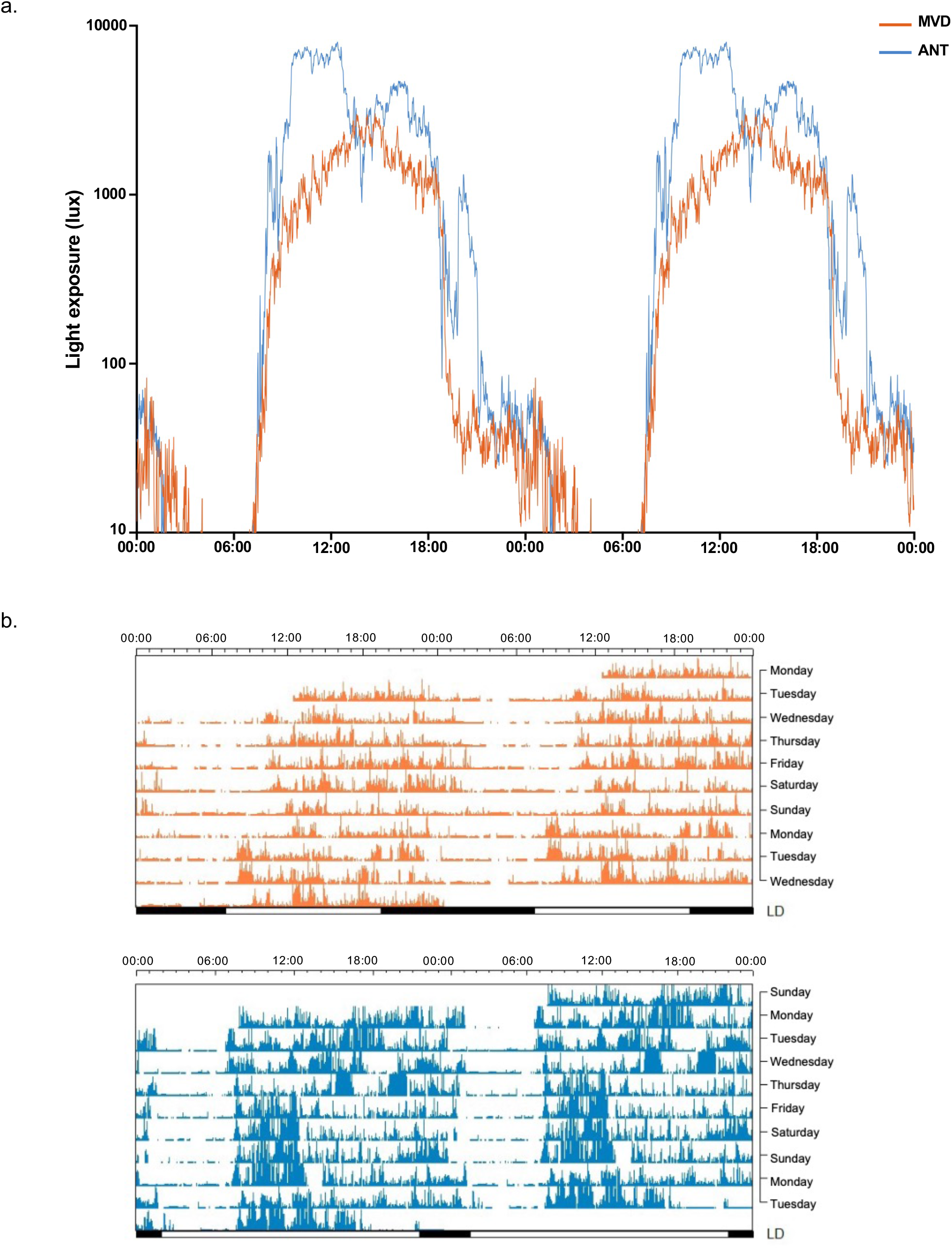
a. Average light exposure (in lux) plotted on a log scale for both locations: Montevideo (MVD, orange) and Antarctica (ANT, blue). Data are double plotted so that light levels across midnight can be more easily observed. b. Representative activity actograms for one participant during the 10 consecutive days of recording for both locations: MVD (top, orange) and ANT (below, blue). Data are presented in rows of 48 h, bars at the bottom indicate light-dark cycle, scale on top indicates time in hours.

Representative activity actograms during 10 consecutive days in both MVD (8 workdays and 2 weekend days) and ANT (all considered workdays) illustrate additional impacts of the Antarctic summer (Fig. 2b; Table 1; Supplementary Fig. S1). First, all participants showed circadian rhythms of activity in both recording sites but with a different timing between them, as the acrophase and the midpoint of the most active 10h period (M10c) were significantly earlier in ANT than in MVD (Table 1). In contrast, the mid-point of the least active 5h period (L5c) did not change between MVD and ANT (Table 1), though the onset of the rest phase was more regular in ANT than MVD across days (Fig. 2b) and among participants (Supplementary Fig. S1). Second, the activity period was longer (more minutes in moderate-vigorous physical activity, MVPA >100 mg, Table 1) and more intense (higher average activity of the most active 10h period, M10). The rest was shorter and quieter (lower average activity of the least active 5h period, L5) in ANT than in MVD. This resulted in a more robust circadian rhythm of activity (higher relative amplitude, RA in ANT with respect to MVD (Table 1). Third, because of the tight schedule shared by all participants during ANT, all phase variables of both activity and light exposure rhythms were less dispersed in ANT than in MVD (Table 1). In line with actigraphy, daily self-reported sleep data showed that sleep was shorter and earlier in ANT than in MVD (Table 1). Interestingly, despite the reduction in sleep duration in ANT, self-reported sleep quality from a subjective visual analogue scale (VAS) included in sleep logs, was not different between ANT and MVD (Table 1).

L5c was strongly correlated with the self-reported mid-sleep point (estimated by sleep logs; R = 0.87, p = 0.001) in MVD as expected, but this association became weaker in ANT (L5c versus mid-sleep point, R = 0.56, p = 0.072). In contrast, the expected association between self-reported sleep quality and duration was observed in ANT (R = 0.73, p = 0.011) but not in MVD (p = 0.11); as well as the association between self-reported sleep quality and RA (ANT: R= 0.70, p = 0.017; MVD: p = 0.1).

### The impact of the Antarctic summer on DLMO and its determinants

Circadian timing was determined by calculating the dim light melatonin onset (DLMO) as marker of the individual circadian phase and, in particular, as indicator of the beginning of the biological night. As previously reported^33^ and shown in Fig. 3a, the DLMO was not significantly different between MVD (22:16 ± 1:14) and ANT (22:01 ± 0:43, Wilcoxon signed-rank test, p = 0.82), although less dispersed in ANT than in MVD. In line with this, as shown in Table 1 and Fig. 3a, L5c, as marker of the individual timing of the rest period was also not significantly different between ANT and MVD (t test, p = 0.11) and less dispersed in ANT. Interestingly, the individual circadian phase shift between ANT and MVD (ΔDLMO), was positively correlated with the individual shift in L5c between both conditions (ΔL5c) (R = 0.69, p = 0.018, Fig 3b).

**Figure 3.**
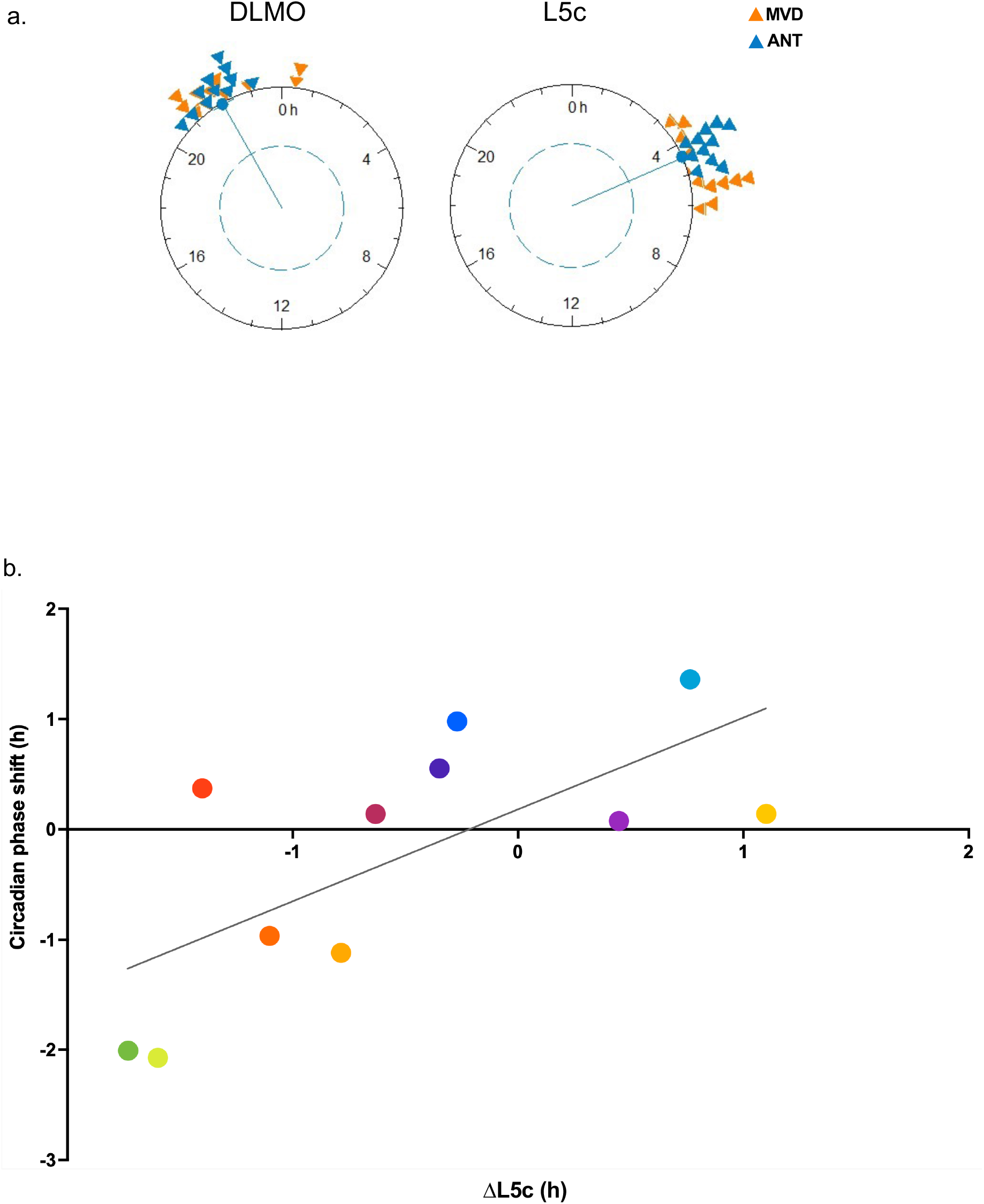
a. Rayleigh’s test for dim light melatonin onset (DLMO) and for the midpoint of the 5 hours of less activity (L5c) for both locations: Montevideo (MVD) in orange, Antarctica (ANT) in blue. b. Pearson linear correlation for the individual circadian phase shift (ΔDLMO) between ANT and MVD with the individual shift in L5c (ΔL5c) between ANT and MVD (R = 0.69, p = 0.018). Each participant is represented with a different color (colors are consistent with Fig. 4b).

The Antarctic summer means a significant increase of light exposure. As shown in Fig. 4a, when considering the individual 3h-phase-advance window with respect to individual DLMO (PAW, 3 h before the DLMO antipode) participants were in average significantly more exposed to light in ANT with respect to MVD (MVD 413.98 ± 155.85 lux vs ANT 1250.80 ± 999.98 lux, Wilcoxon signed-rank test p = 0.019). Likewise, when considering the individual 3h-phase-delay window (PDW, 3 h before the DLMO), participants were in average significantly more exposed to light in ANT with respect to MVD: (MVD 50.09 ± 39.06 lux vs ANT 346.18 ± 151.27, Wilcoxon signed-rank test p = 0.001). This great increase in light exposure in both light-sensitive windows had overall opposite effects on the circadian phase, and this made it necessary to analyze individual differences in light exposure between MVD and ANT. The individual circadian phase shift between ANT and MVD (ΔDLMO) was negatively correlated with the individual difference in light exposure in the PAW (R = -0,80, p = 0.003, Fig 4b) but was positively correlated with the individual difference in light exposure in the PDW (R = 0.747, p = 0.008, Fig. 4b). In the PAW the greater the light exposure in ANT with respect to MVD (high Δlight exposure in the PAW), the earlier the DLMO (negative ΔDLMO) in ANT with respect to MVD. Moreover, participants who showed very low difference or who were even exposed to less light in ANT than in MVD (Δlight exposure in the PAW around zero) exhibited a circadian phase delay (positive ΔDLMO). In contrast, in the PDW the greater the light exposure in ANT with respect to MVD (high Δlight exposure in the PDW), the later the DLMO (positive ΔDLMO) in ANT with respect to MVD. Accordingly, participants who showed little or no difference in light exposure in ANT than in MVD (low Δlight exposure in the PDW) exhibited a circadian phase advance (negative ΔDLMO).

**Figure 4.**
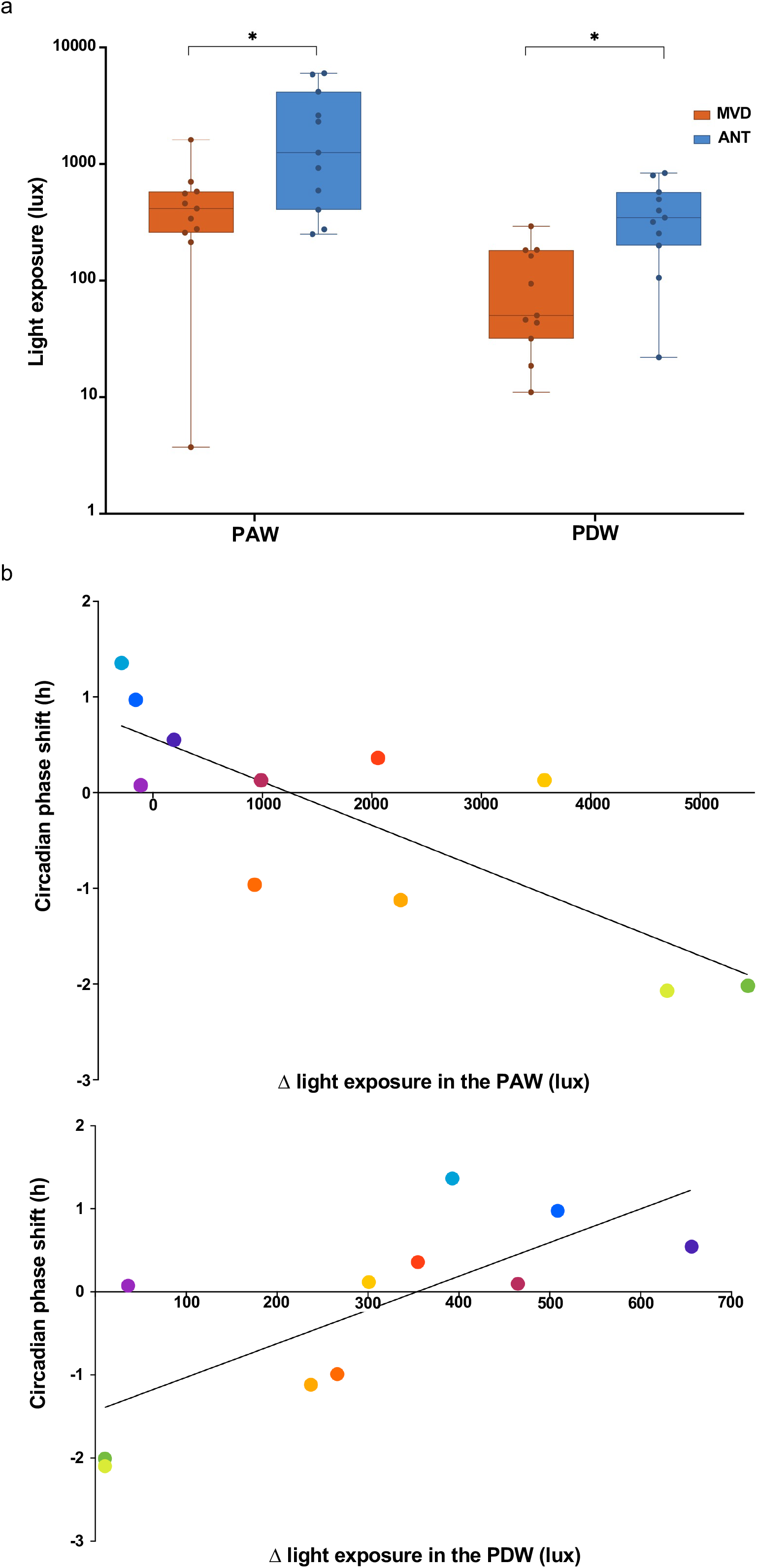
a. Light exposure (lux) plotted on a log scale for the phase advance window (PAW) and phase delay window (PDW) in both locations: Montevideo (MVD, orange) and Antarctica (ANT, blue). Light exposure was different in both the PAW (Wilcoxon singed-rank test, p = 0.019) and the PDW (Wilcoxon singed-tank test, p = 0.001).b. Pearson linear correlation for the individual circadian phase shift (ΔDLMO) between ANT and MVD with the difference in light exposure in the PAW (on top, R = -0.8, p = 0.003), and in the PDW (below, R = 0.75, p = 0.008) between ANT and MVD. Each participant is represented with a different color (colors are consistent with Fig. 3b).

## DISCUSSION

Although underexplored experimentally, trans-latitudinal trips are postulated to have impacts on the circadian system when they mimic acute seasonal changes in photoperiod^40^. In this study, the great amount of light participants were exposed to in the Antarctic summer impacted on the activity-rest rhythms, sleep patterns, and circadian phase of Uruguayan university students. During their normal lives around the equinox in Montevideo, Uruguay, the light exposure participants were used to was different among them and these differences were key to evaluate the effects of the Antarctic summer. Overall, the activity-rest rhythm was earlier, more robust, and less variable among participants in ANT with respect to MVD, while self-reported sleep was also earlier and shorter. More importantly, the shifts of the circadian phase and sleep patterns observed in ANT were driven by participants’ individual habits in MVD: a) participants who advanced their sleep timing in ANT respect to MVD, also advanced their DLMO in ANT, while participants who delayed their sleep timing in ANT respect to MVD, also delayed their DLMO in ANT; and b) shifts of the circadian phase depended on the individual light history; i.e., DLMO was advanced in participants exposed to more light in the phase-advance window in ANT, and delayed in participants exposed to more light in the phase-delay window in ANT.

The participants of this study, who were used to a normal individual lifestyle in MVD, including personal specific profiles of light exposure and social demands, were subjected to a 10 days-summer trip to an Antarctic scientific base, in which they shared a daily routine of classes, field work, meals, social activities, and conditioned sleep schedules^33,45^. Unlike previous studies that mainly focused on the impact of the huge increase of light in the Antarctic summer^46,47^, our experiment aimed to evaluate the effect of individual differences in light exposure between the two conditions. The rationale of this ecological chronobiology experiment relied on that the baseline situation (MVD) was not a fully controlled condition. Participants in MVD were not indicated to follow any protocol of either scheduled activity or exposure to light; rather they just displayed their normal lives and thus had variable patterns of activity and light exposure. In contrast, the Antarctic summer was a homogeneous situation among participants, who shared the same strict schedule of activities imposed by the Antarctic scientific base agenda. The social synchronization generated by the schedule of scientific bases in Polar regions is indisputable as shown by previous self-reported and objective data from short summer trips^33,45,47,48^. In the case of this study, the social synchronization imposed by ANT was crucial for the emergence of striking differences in individual light exposure between the two conditions; and consequently, it enabled us to assess the effect of individual habits, specially of light history.

Objective reported parameters of activity patterns (Table 1) were consistently earlier in ANT than in MVD (except for L5c that did not change), but in contrast to an Arctic summer trip study that showed a delay in the acrophase^48^. In addition, although all these objective parameters were more variable among participants in MVD, a set of associations revealed that the individual circadian system was in a stable balanced state in this baseline-life condition. For example, parameters that are expected to correlate as comparable indicators provided by different instruments (L5c from actimetry, self-reported midsleep point, and DLMO) were significantly associated in MVD. The loss of these expected associations in ANT suggests that the Antarctic summer induced a misalignment of the circadian system.

This vision of the suffering of the circadian system in ANT is in line with previous studies that have argued that the extreme light condition of the Antarctic summer acts as an apparent stressor to the circadian clock, decreasing the quality of sleep, delaying melatonin phase, and causing desynchronization^38,46,47,49^.

Sleep quality is vital to individual health and wellbeing and involves individual’s self-satisfaction with all aspects of the sleep experience^50^. Sleep quality is therefore associated with longer sleep duration, proper timing, high efficiency, and less fragmented sleep^51–56^. Participants’ sleep quality, estimated by VAS, did not change between MVD and ANT in line with one of the few studies reporting sleep changes in a summer Antarctic expedition^57^. This lack of change in sleep quality probably depended on the interaction of multiple causes acting in opposite directions. On one hand, self-reported sleep duration in ANT was less than the minimum of 7 h recommended for young adults (Table 1)^45^, with potential detrimental effects on sleep quality. On the other hand, actimetry recordings showed that sleep was less fragmented in ANT as the activity-rest rhythm was more robust in ANT than in MVD (RA in Table 1). Interestingly, these contradictory influences resulted in the finding of the expected associations between sleep quality and both sleep duration and fragmentation in ANT but not in MVD. In line with this, a similar population of Uruguayan university students, was already reported to have low sleep quality during their normal lives that also did not correlate with sleep duration (Paz, V. personal communication).

DLMO is an accurate, non-invasive, and reliable measure of the endogenous circadian rhythm^13,14^, whose shifts are also used to show variations of the circadian phase to adapt to acute transitions^58^ even in real-life situations^16,20,33^. Although it is well known that bright light is the main cue that modulates the body clock timing^23^, it has been reported that age^24,59,60^, chronotype^61,62^, and social demands (e.g., school or work schedules)^63–65^, among others, interact with light in the modulation of human circadian phase inducing a wide individual variation. Interestingly the lack of an average DLMO shift between MVD and ANT that can be interpreted at first glance as the absence of effect, is indeed the demonstration of how much individual attributes matter^33^. Although neither DLMO nor L5c changed in average between MVD and ANT, participants’ individual circadian phase shifted importantly between both conditions in a comprehensive way that depended on both their circadian preferences^33^ and their sleep patterns (Fig. 3). Moreover, the circadian phase and sleep timing changed in concert: a) both DLMO and L5c were more dispersed in MVD than in ANT as an additional evidence of the social synchronization the Antarctic summer imposed; b) both shifts in DLMO and L5c were associated showing a predictable effect of the change of sleep timing on the shift of the circadian phase, and c) the magnitude of both shifts was comparable; i.e., when sleep was advanced by 2 h, the DLMO accompanied this change with a 2-h advance. On the other hand, the strong association between these two different objective phase parameters (DLMO and L5c) proves the reliability of our data by showing that each individual reacts in a holistic way.

As the light-dark cycle is the most important entrainer of the circadian system across evolution^66,67^, the way that light influences the DLMO has been extensively documented in controlled experimental conditions; namely DLMO can either be advanced by bright light exposure in the morning^5,68–71^, or delayed by light exposure in the evening^24,72^. The precise moment these light sensitive PAW and PDW occur is, of course, dependent on the individual DLMO and should thus be calculated considering the individual circadian time^5,73^. In this study, light exposure was significantly higher in average in ANT with respect to MVD in both the PAW and the PDW, as expected (Fig. 4a). However, the difference in light exposure between ANT and MVD in these individual light-sensitive windows was less pronounced than the difference observed during the morning and the evening (Fig. 2a). Indeed, individual changes in light exposure between ANT and MVD were highly variable in both the PAW and the PDW, and even negative for three participants in the PAW (Fig. 4b). Interestingly, the two participants who showed the greatest difference in light exposure in the PAW were the ones who showed the least change in the PDW (Fig. 4b). These results confirm the importance of the individual assessment not only of the light-sensitive windows, but also of the evaluation of individual differences between conditions in line with the current debate of the importance of individual photic history^21–23^.

Long-term photic history shapes the individual physiological response to light and underlies plastic changes of the circadian phase across seasons and less gradual transitions such as trans-latitudinal trips or between urban and natural environments^16,20,23,40^. Although it is clear that photic history will affect subsequent response to light, only a few studies have focused on light exposure differences to assess phase shifts in real-life conditions^38,74^. Transitions between locations with artificial to natural lighting induce acute phase-advance shifts attributable to a concerted change in light exposure, which is simultaneously increased in the PAW and decreased in the PDW^16,20^. In contrast, as any seasonal change towards the summer, the Antarctic trip means an increase of light exposure in both light sensitive windows with opposing effects on the circadian phase, and therefore with a theoretical unpredictable outcome. We experimentally confirmed this impossibility of predicting the valence of the circadian phase shift. When light exposure increased at the same time in both the PAW and the PDW, the average DLMO did not change between ANT and MVD. However, our study also highlights the need to focus on individual light history as the higher the difference in light exposure in the PAW between ANT and MVD, the greater the advance of their DLMO; and vice versa, the higher the difference in light exposure in the PDW between ANT and MVD, the greater the delay of their DLMO (Fig. 4b). Therefore, this study allowed us to emphasize the importance of individual light history as a proxy of circadian phase shifts because: a) the changes in light exposure in the PAW and the PDW between ANT and MVD were not homogeneous among participants (Fig. 4b); and b) the light exposure change between ANT and MVD exhibited by the majority of participants was specific to one of the sensitive windows (PAW or PDW), thus determining the valence of the individual circadian phase shift. This is the first study to show a clear physiological effect of light either advancing or delaying the circadian phase dependent on individual light history in an ecological study.

## Limitations

We recognize several limitations of this study, most of them derived from the fact that this is an ecological study. First, the reduced number of participants from a homogeneous population of university students prevents the generalization of the interpretation of our results. Second, the Uruguayan Summer School on Introduction to Antarctic Research as a training course, in each edition carefully selects 12 undergraduate science students to travel to Antarctica. Thus, the volunteer participants in this study were not randomly selected and were highly motivated to be part of this study. Third, although all the procedures were personally supervised by the authors of this study, especially during ANT, light information provided by wrist actimeters could be underestimated because they may have eventually been covered by warm clothing (jackets and gloves). Fourth, because the impact of ANT was contrasted with normal life (MVD), we did not control for other stimuli that could affect the circadian phase (food intake, stimulants, substance use, stress, etc.). However, we do not consider that it has been a limitation of this study not having had a completely controlled situation of participants’ life in MVD, as this allowed individual differences to emerge, which are the focus of this study.

### Concluding remarks

Our understanding of the complex interactions between photic and non-photic stimuli for the daily adjustment of the human circadian phase has made important progress in the last decade. First conceived as a rather stable individual trait, strong evidence has shown that, for example, the circadian phase can be very plastic^65^, light sensitivity is highly diverse among individuals^11^, and the power of light entrainment can be strong and fast even in ecological conditions^16,20^. People living in the same city (or even in the same house) can be subjected to very different light exposure patterns related to personal school/work schedules, chronotypes, and lifestyles. In this study, we show the importance of this individual light history when it is challenged by a sudden and synchronized increase in light exposure affecting both light-sensitive circadian phase windows. It is not the absolute amount of light that all the participants received in a similar way in Antarctica what matters to explain the shifts of the circadian phase induced by this short trip to the Antarctic summer. Rather, the ultimate cause of each participant’s distinctive circadian phase shift relied on the unique change in light exposure each individual was subjected to between his/her baseline life in Montevideo and the Antarctic summer. We show that both light-sensitive windows were similarly capable of influencing the net circadian phase shift; i.e., the higher the light exposure difference in the PAW or in the PDW the higher the circadian phase advance or delay, respectively. This remarkable finding is supported by the diverse individual light history among participants: those participants whose difference in light exposure was similar in both the PAW and the PDW did not show a significant circadian phase shift, while those participants whose difference in light exposure was prevalent in the PAW advanced their circadian phase and those participants whose difference in light exposure was prevalent in the PDW delayed it.

This ecological study not only contributes solid evidence on the importance of light history as predictor of circadian phase shifts, but also reinforces the need to pay attention to individual photic history in future studies interested in the modulation of the human circadian phase, especially in normal-life conditions.

## METHODS

### Participants

Eleven healthy students from Facultad de Ciencias, Universidad de la República, Uruguay participated in this study out of the twelve members of the Second Uruguayan Summer School on Introduction to Antarctic Research held in January 2016, in the Uruguayan Antarctic Scientific Base Artigas. All participants were clinically assessed to confirm they were healthy, and none showed sleep disturbances or depression signs (Beck’s Depression Inventory score < 10 ^75^. All procedures were approved by the ethics committee at the Instituto de Investigaciones Biológicas Clemente Estable, Ministerio de Educación y Cultura, Uruguay (CEP/HCPA 14-0057; approval date: December 16, 2013) and complied with the principles outlined by the Declaration of Helsinki^75^. All participants signed an informed consent form stating that they had been told about the aims and procedures of the study, their right to end participation without any explanatory statement at any time, their data being coded as to maintain anonymity, and their data being communicated for scientific purposes only.

We compared two different conditions: baseline situation, which corresponded to the start of university semester in the austral fall equinox 2016 (March 7–17, 2016, Montevideo, Uruguay, 34°540 S; 56°110 W, LD 12:12) and the Antarctic trip, which corresponded to the 2016 Uruguayan Summer Antarctic School (January 17–27, 2016, King George Island, 62°110 S; 58°520 W, LD 20:4). In the Montevideo baseline situation (MVD), students developed their regular lives, attending university courses with no restrictions nor special instructions. During the Antarctic trip (ANT), all the participants were living together, followed the same daily agenda (mealtime, type of food intake, activities, etc.), and had an intense work schedule that was carried out regardless of the day of the week (workdays or weekend). Therefore, in MVD sampled weekends were excluded and only workdays were analyzed.

### Chronotype

Chronotypes were assessed using the Spanish versions of both the Munich Chronotype Questionnaire (MCTQ)^41,42^ and the Morningness - Eveningness Questionnaire (MEQ)^43,44^. These questionnaires were answered during the first day of the MVD. MCTQ reports were used to calculate the mid-sleep point on free days corrected for sleep debt on work days (MSFsc)^32^ as an indicator of individual chronotype, in which higher values indicate later chronotypes. MEQ scores were calculated directly from the answers about sleep/activity preferences, and in this case, higher scores indicate earlier circadian preferences^44^.

### Sleep logs (SL)

Participants were instructed to fill sleep logs with questions about the previous night, every morning during sample periods on both conditions. From the self-reported time of sleep onset and sleep end we calculated sleep duration (time interval from sleep onset to sleep end) and mid-sleep point (midpoint between sleep onset and sleep end). We also estimated sleep quality from a subjective visual analogue scale (VAS) extending from “very bad night” to “very good night”^76^.

### Melatonin

Hourly saliva samples (18:00 - 00:00 h) were collected in dim light (< 30 lux) on the last night of each sample period. Following Lewy^13,14^, saliva samples were obtained in the same settings for all participants in both conditions; i.e. participants remained in resting position in dark rooms, except for brief trips to the also dark bathroom from 17:00 to 00:00. Samples were then frozen and stored at -20°C, and later assayed for melatonin using the salivary melatonin competitive ELISA kit from Salimetrics following the manufacturer’s instructions. Tests were carried out by the same technician at the same time for 3 consecutive days, and control and treatment samples were combined in each ELISA plate. The concentration-response curves obtained were indistinguishable from those reported by the manufacturer. The same applied to the IC10 values (melatonin concentrations inhibiting 10% of the absorbance signal observed in the absence of melatonin), taken as an indication of assay sensitivity. This validates the use of the analytical sensitivity value reported by the manufacturer (1.37 pg/mL melatonin).

Circadian timing was determined by calculating the dim light melatonin onset (DLMO)^14,77,78^, as a marker of the individual circadian phase and, in particular, as an indicator of the beginning of the biological night. Following Silva et al.^33^ and Coirolo et al.^73^, individual basal melatonin levels were calculated as the average of the melatonin levels before the nocturnal increase. Individual DLMO was calculated as the interpolated point in time at which the quadratic fit surpassed 2 SD above the individual basal melatonin level in MVD samples. For each participant, the melatonin level corresponding to MVD DLMO was considered the individual melatonin threshold. ANT DLMO was calculated as the time at which the melatonin level reached the individual threshold. To measure the individual impact of ANT on the circadian phase we calculated the circadian phase shift as the difference between DLMO values for each participant: ΔDLMO = DLMO_ANT_ – DLMO_MVD_.

### Actigraphy

Participants wore ambulatory wrist activity and light monitors on the non-dominant arm (GENEactiv Originals, Activinsights) continuously during both sample periods. The device incorporated an expandable wristband allowing use over outdoor clothing. All devices were configured on the same computer and set to record at 10 Hz. Data was exported and converted to 1-min epochs using GENEActiv Windows Software from Activinsights. We analyzed actigraphy data with the integrated software El Temps (© Antoni Díez-Noguera, Barcelona, CA, Spain). From the activity and light exposure, we obtained individual actograms for all participants in both conditions. We also performed cosinor and non-parametric analyses on both physical activity and average light exposure data (in lux). These analyses yielded linear variables, including the average activity of the most active 10h period (M10), the average light exposure of the brightest 10h period (MB10), the average activity of the least active 5h period (L5) and the average light exposure of the least bright 5h period (LB5)^79,80^. We then estimated the activity relative amplitude (RA) as M10-L5/M10+L5^80,81^. We also estimated phase variables, including acrophase (the moment of the day in which the cosinor function reaches its maximum for both physical activity and light exposure); M10c (the midpoint of the most active 10h period); MB10c (the midpoint of the brightest 10h period); L5c (the midpoint of the least active 5h period); and LB5c (the midpoint of the least bright 5h period)^19^. For physical activity, the raw 10-Hz triaxial data were first summarized into the gravity-subtracted sum of vector magnitudes for each 1-min epoch intervals. The resulting units for this outcome variable are g·min^82,83^. We then calculated the average gravity-subtracted signal vector magnitude for each minute, thus converting physical activity values from a time-dependent unit (g·min) to the time-independent milligravitational units (mg) in order to compare with previously reported cut-points (moderate-vigorous physical activity above 100 mg, MVPA)^84,85^. To measure the impact of sleep timing as determinant of the change of individual circadian phase between ANT and MVD, we calculated the difference between L5c values for each participant: ΔL5c = L5c_ANT_ – L5c_MVD_.

We calculated for each participant the hourly average of light exposure across each sample period in both conditions in two 3h-phase-relevant windows centered by individual DLMO ^5,73^: in the phase-delay window (PDW, 3 h before the DLMO) and in the phase-advance window (PAW, 3 h before the DLMO antipode). To measure the impact of light as determinant of the change of individual circadian phase between ANT and MVD, we calculated the difference between the average light exposure in both the PDW (Δ light exposure PDW= light exposure PDW_ANT_ - light exposure PDW_MVD_) and the PAW (Δ light exposure PAW= light exposure PAW_ANT_ - light exposure PAW_MVD_) for each participant.

### Statistical analysis

Data are expressed as mean values ± standard deviation throughout the text unless otherwise stated. As data did not comply with normality and/or homoscedasticity, statistical comparisons were analyzed by non-parametric test (Wilcoxon signed-rank test) for paired-samples comparison between ANT and MVD. Associations between variables were assessed by Pearson correlations. Statistical procedures were carried out the software PAST^86^. Values of p ≤ 0.05 were considered statistically significant.

## Acknowledgements

We wish to thank the students of the Second Summer School on Introduction to Antarctic Research, who kindly volunteered as participants of this study. We are especially thankful to the Instituto Antártico Uruguayo for the logistic support. Financial support was provided by Comisión Sectorial de Investigación Científica (CSIC I+D_2016/623 and CSIC Grupos 2018 #92) and Facultad de Ciencias, Universidad de la República, Montevideo, Uruguay. Julieta Castillo was supported by a scholarship from Agencia Nacional de Investigación e Innovación, Uruguay.

## Date availability

All data reported in this paper will be shared by the corresponding author upon request. Any additional information required to reanalyze the data reported in this paper is available from the corresponding author upon request.

## Authors contribution

Funding, project administration, experimental design, interpretation of data and writing original draft, BT and AS.; data processing, statistical analysis and figures design, JC; supervision of actimetric data processing and review of the manuscript ACT and MPH. All authors contribute to the writing of the manuscript.

